# Regulation of translational fidelity by repression of C/D box SNoRNPs during UPR

**DOI:** 10.1101/2025.05.09.653119

**Authors:** Wenyuan Zhao, Ananya Gupta, Sanjeev Gupta

## Abstract

Endoplasmic reticulum (ER) is the major cellular compartment where the folding and maturation of secretory and membrane proteins take place. When protein folding demands exceed the protein handling capacity of ER, the unfolded protein response (UPR) pathway is activated. UPR reduces the client protein load in the ER by inhibiting protein translation and increases protein handling capacity by upregulating genes encoding ER-resident molecular chaperones. The main pathway for translational repression in response to ER stress has been the phosphorylation of eIF2α by PERK, which also suppresses rRNA synthesis. rRNA methylation and pseudouridylation can fine-tune protein synthesis. Small nucleolar ribonucleoproteins (SnoRNPs) are mainly separated into two subtypes: C/D box and H/ACA box. C/D box SNoRNPs guides rRNA 2’-O-methylation, while H/ACA SNoRNPs catalyze the pseudouridylation modifications of rRNAs. We hypothesize that the UPR may regulate translational fidelity by regulating the expression of core proteins of C/D and H/ACA box SnoRNPs. In this study, we show that the expressions of NHP2L1, NOP56 and FBL were significantly downregulated by UPR. In addition, FBL knockdown reduced cell proliferation and colony formation. Furthermore, we show the effects of UPR and FBL knockdown on rRNA methylation and translation fidelity, including alterations in nonsense suppression, frameshifts, ribosome pausing and translation initiation by IRES. In conclusion, this study reveals the regulation of translational fidelity during UPR by change in rRNA methylation due to reduced function of C/D box SnoRNPs.

## Introduction

The endoplasmic reticulum (ER) serves as the primary cellular organelle for the proper folding and maturation of secretory and membrane proteins [1]. When the protein folding demand exceeds the ER’s capacity, ER stress occurs, thereby activating a series of cellular complementary adaptive responses known as the unfolded protein response (UPR) [2]. UPR involves an intricate network of signaling pathways originating from three EnR transmembrane sensors: ATF6, PERK and IRE1 [3, 4]. The UPR pathways modulate gene expression to increase ER’s protein handling capacity and suppresses protein translation to restore ER homeostasis [2, 5]. Specifically, the activation of PERK during ER stress leads to the phosphorylation of eIF2α, which attenuates protein synthesis by inhibiting cap-dependent translation initiation [6-9]. In addition, PERK-dependent phosphorylation of eIF2α plays a critical role in the inactivation of RRN3/TIF-IA, leading to the downregulation of ribosomal RNA (rRNA) synthesis [10].

Small nucleolar RNAs (SnoRNAs) are a family of small non-coding RNAs located in the nucleolus and guide the post-transcriptional modifications of rRNAs. Small nucleolar ribonucleoprotein (SnoRNP) is a stable and functional assembly of SnoRNA and specific core proteins. It is mainly classified into two subtypes: C/D box and H/ACA box SnoRNPs. C/D box SnoRNP catalyzes the 2’-O-methylation of rRNA, whereas H/ACA box SnoRNP catalyzes the pseudouridylation [11]. The core proteins of C/D box SnoRNP include NHP2L1 (or SNU13), NOP56, NOP58 and FBL. Among them, NHP2L1 binds to the K-turn motifs that position FBL at the methylation site, enabling FBL to perform the methyltransferase function [11]. It was reported that FBL overexpression enhances cell proliferation and drug resistance in MCF7 breast cancer cell line [12].

Ribosomes modulate the translation of mRNAs through targeted interactions with them [13]. These interactions can be affected by ribosome heterogeneity through alterations in rRNA and protein composition [14]. THE rRNA modifications are critical for the proper folding, stability and function of rRNA, thereby affecting protein synthesis [15]. Two primary types of modifications—2’-O-methylation and pseudouridylation—constitute about 95% of rRNA modifications, while the rest are various base modifications [16]. Disruptions in these modifications lead to defective protein synthesis, affecting tumour cell growth and survival.

As a primary rRNA modification, the 2’-O-methylation is guided by C/D box SnoRNP and catalyzed by its core protein FBL [16]. The methylation of rRNA by C/D box SnoRNP is considered to reduce the hydrophilic nature of nucleotides and facilitate rRNA to be buried into the ribosome [11]. This modification regulates the structure and stability of rRNA, thereby affecting its functionality [17]. It was found that rRNA hypomethylation significantly affects the fidelity of protein synthesis [18]. We hypothesize that the UPR may regulate translational fidelity by modulating the expression of core proteins of C/D and H/ACA box SnoRNPs.

Here, we evaluated the expression of core proteins belonging to C/D and H/ACA box SnoRNPs during ER stress. This study also suggests that NHP2L1, NOP56 and FBL were significantly downregulated by UPR. In addition, FBL knockdown reduced cell proliferation and colony formation. We also selected four critical translation regulation pathways for further investigation: ribosome pausing [19, 20], IRES-regulated cap-independent translation [21, 22], ribosome frameshifting [23, 24] and stop codon readthrough (190UGA) [25]. Furthermore, we show the effects of UPR and FBL knockdown on rRNA methylation and translation fidelity, including alterations in nonsense suppression, frameshifts, ribosome pausing and translation initiation by IRES.

## Materials and Methods

### Chemicals and Antibodies

AA147 (# HY-139293) and IXA4 (# HY-139214) were purchased from MedChem Express. CCT020312(# 324879) and GSK2606414(# 516535) were purchased from Merck Millipore. Brefeldin A (BFA; # 1131/5), Thapsigargin (TG; # 1138) and Tunicamycin (TM; # 3516) were purchased from Tocris. The primary antibodies such as β-actin (A5060) were purchased from Sigma-Aldrich. FBL (ab5821) and NOP56 (ab229497) were purchased from Abcam. PERK (C33E10), XBP1s (143F), CHOP (L63F7), eIF2α (9722S) and P-eIF2α (9721S) were purchased from Cell Signaling Technology.

### Cell culture

Cells were maintained in high-glucose Dulbecco’s modified eagle’s medium (DMEM; Sigma-Aldrich, Cat no: D6429) supplemented with sterile-filtered 10% FBS (Sigma-Aldrich, Cat no: F7524), 1% penicillin-streptomycin (P/S; Sigma-Aldrich, Cat no: P0781), 1% L-Glutamine (Sigma-Aldrich, Cat no: G7513) and 1% Na-Pyruvate (Sigma-Aldrich, Cat no: S8636). They were incubated at 37°C in a humidified atmosphere of 5% CO_2_.

### MTS assay

The MTS reagent (ab223881), which contains the tetrazolium inner salt, was used to assess cell viability during proliferation. To prepare the reagent, MTS powder was dissolved in DPBS at a concentration of 2 mg per 1 ml. PMS powder was separately dissolved in DPBS at a concentration of 0.9 mg per 1 ml. Then, 100 μl of the PMS solution was added to 1 ml of the MTS solution and thoroughly mixed. Next, 20 μl of the resulting MTS solution was added to each well and incubated at 37°C for 1 h. The conversion was measured at an absorbance of 490 nm using a microplate reader, with medium-only wells serving as blanks. For example, parental MCF7 cells were seeded at 2000 cells per well in a 96-well plate (five replicate wells per set) on day 0 with a final volume of 100 μl per well. Four plates were prepared for taking measurements on days 1, 2, 3 and 4.

### Colony formation assay

Taking parental MCF7 cells as an example, 500, 1000 or 2000 cells per well were seeded in a 6-well plate containing 2ml of complete DMEM. The plate was then placed in a humidified incubator at 37°C for 14 days. After incubation, the wells were washed twice with DPBS, and the cells were fixed with 4% formalin for 30 min. Next, the cells were stained with 0.1% crystal violet for 20 min to visualize the clones. The wells were subsequently washed with DPBS, allowed to dry, and the number of clones was counted using ImageJ.

### Quantitative real-time polymerase chain reaction (qPCR) qPCR procedure

Total RNA was isolated using Trizol reagent (ThermoFisher 15596026), chloroform (Sigma-Aldrich 372978), isopropanol (Fisher Scientific 11388461) and 75% ethanol (Fisher Scientific 16695992). cDNA synthesis was performed using the HiScript II 1st Strand cDNA Synthesis Kit (R211-02). qPCR was conducted using the ThermoFisher kit (Cat no: 4440040) on an Applied Biosystems 7500 Real-Time PCR machine, following the manufacturer’s two-step amplification protocol: an initial step at 50°C for 2 min and 95°C for 10 min, then 40 cycles of 95°C for 15 sec and 60°C for 1 min. Data were analyzed using the 2^-ΔΔCt^ method. Primers used in qPCR are listed in S1 Table.

#### Assessment of rRNA methylation levels using qPCR

Belin *et al*. were the first to use the qPCR technique to measure rRNA methylation levels by performing qPCR at both low and high dNTP concentrations [12]. Under high dNTP conditions, rRNA samples yield full-length cDNA after reverse transcription. However, in methylated samples, reverse transcription is interrupted at methylation sites under low dNTP concentrations, reducing the yield of full-length cDNA. This process enables the assessment of methylation levels, resulting in higher CT values for cDNA synthesized at low dNTP concentrations compared to those at high concentrations. Therefore, we used the qPCR technique to assess the methylation level of the 2’-O-methylated sites in the target RNA samples.

The qPCR results were analyzed using two different calculation methods [26, 27]. The first method [26] used the 2^(CTlow -CThigh)^ function to assess the methylation level. The fold change in rRNA methylation was then calculated by comparing samples with reference gene and target genes. The second method [27] assessed the methylation level using RT efficiency. First, the average CT values were calculated from duplicate measurement for each sample, condition and target gene. Then the delta CT (ΔCT) was calculated. For reactions with high dNTP concentrations, ΔCT was determined by subtracting the average CT value of the reference gene from the average CT value of the target gene for each condition. For low dNTP reactions, ΔCT was calculated by subtracting the CT value of each shFBL sample from the CT value of the scramble control. Next, the delta-delta CT (ΔΔCT) for samples with high dNTP concentrations was calculated by subtracting each ΔCT from the average ΔCT of the scramble control. Then the relative quantification (RQ) value for each sample with target genes were determined using the functions 2^ΔΔCT^ and 2^ΔCT^ for high dNTP and low dNTP, respectively. Finally, the RT efficiency was determined by dividing the RQ value of the low dNTP samples by the corresponding RQ of the same samples under high dNTP concentrations.

### Western blotting analysis

Total cell protein samples were loaded onto SDS-PAGE gels and run at 110 V for 90 min, until the bromophenol blue (loading dye) reached the bottom of the gel. Proteins were then transferred to a nitrocellulose membrane (Sigma-Aldrich GE10600001) through semi-dry transfer. After transfer, the membrane was blocked for 2 h at room temperature with milk or BSA in PBST or TBST, depending on different antibodies in S2 Table, followed by an overnight incubation at 4°C with the primary antibody. The next day, the membrane was washed three times with PBST or TBT, then incubated with the secondary antibody for 2 h at room temperature. The proteins were detected using Western Lightning Plus-ECL (PerkinElmer NEL104001EA), and β-actin was used as the control.

### Lentivirus preparation

On day 1, 3.5 million 293T cells were seeded in a T75 flask containing complete DMEM. On day 2, 10.5 µg of total plasmid (3.5 µg psPAX2, 1.75 µg pMD2G, and 5.25 µg lentiviral transfer plasmid) were transfected using JetPEI solution (Polyplus Transfection 101-10N). After 16 h, the medium was replaced with fresh complete medium containing 4 mM caffeine. On day 4, the supernatant containing lentivirus vectors was harvested, filtered through a 0.45 µM filter and stored at -80°C.

### Reporter assay

#### Firefly/Renilla dual luciferase reporter assay

Taking MCF7 parental cells as an example, 0.15 million cells were seeded in a 6-well plate on day 1. On day 2, 800 ng of the target plasmid was transfected using Turbofect (Thermo Scientific R0531). Cell lysates were harvested on day 3, and Firefly/Renilla dual luciferase activities were assessed according to the manual of Millipore Cat. SCT152. Signal calculations were performed as (Firefly signal - blank) / (Renilla signal - blank).

#### β-Galactosidase based reporter assay

Taking MCF7 parental cells as an example, 0.15 million cells were seeded into 6-well plates on day 1. On day 2, 800 ng of the target plasmid and 200 ng of the β-Gal plasmid were transfected using Turbofect (Thermo Scientific R0531). On day 3, cell lysates were harvested on day 3, and β-Galactosidase activity was assessed in a 96-well plate by adding 50 μl of lysate and 50 μl of Assay 2X to each well (in triplicate). The plate was incubated at 37°C for 30 min, or until a faint yellow colour appeared, not exceeding 3 h. To stop the reaction, 150 µl of 1M Sodium Carbonate was added to each well. Absorbance was measured at 420 nm.

## Results

### Regulation of C/D box SnoRNPs by UPR

The parental MCF7 and T47D cells were treated with BFA and TG for 24 h and expression of SnoRNP core proteins was quantified by qPCR. Fig 1A shows that NOP56 expression was significantly downregulated after TG and BFA treatments in parental MCF7 cells. The expressions of NHP2L1 and FBL were downregulated after TG treatment only, while NOP58 expression did not change. Fig 1B shows that expressions of H/ACA box SnoRNP core proteins exhibited no significant differences following treatment with TG or BFA, except for GAR1, which was upregulated after TG treatment. Fig 1C shows that the expressions of all C/D box SnoRNP core proteins were significantly downregulated after TG treatment in parental T47D cells. Moreover, NHP2L1 and NOP56 were downregulated after BFA treatment, while NOP58 and FBL did not change. Fig 1D shows that NOP10 and DKC1 were downregulated after BFA and TG treatments, respectively, while GAR1 was significantly upregulated by BFA. Therefore, the expressions of FBL, NHP2L1 and NOP56 were significantly decreased following TG treatment in both MCF7 and T47D cells, which are members of C/D SnoRNP complex.

**Fig 1.**
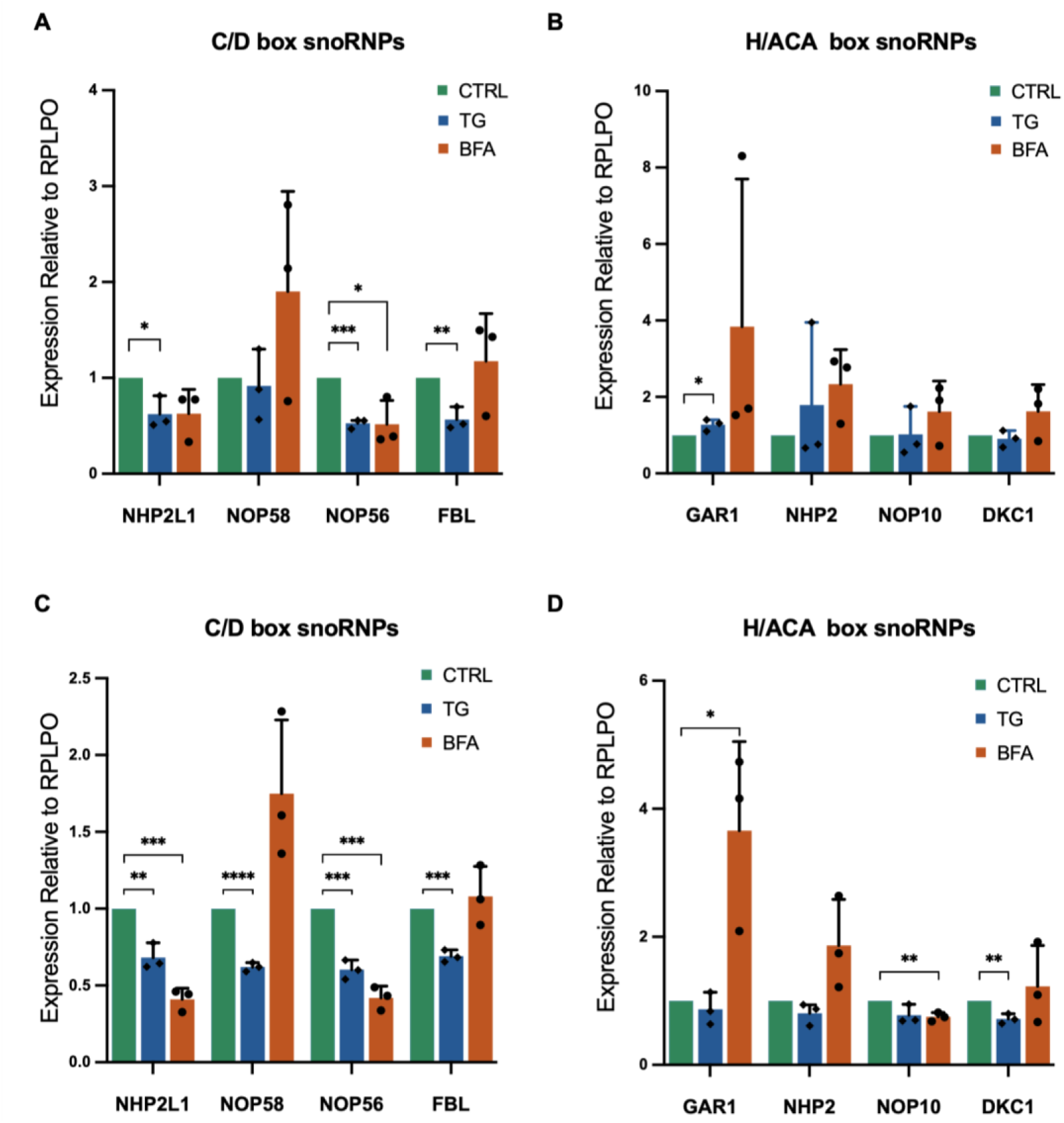
Expression of SnoRNP core proteins during UPR. (A, B) MCF7 cells and (C, D) T47D cells were treated with BFA (0.5 μg/ml) and TG (1 μM) for 24 h. The expressions of (A, C) C/D box and (B, D) H/ACA box SnoRNP core proteins were quantified by qPCR (n=3), normalized to RPLPO. Error bars represent mean ±S.D. from the three independent experiments. Statistical analysis was performed using unpaired t-tests. *P<0.05, **P<0.01, ***P<0.001, ****p<0.0001, and n.s. denotes non-significant results.

To investigate the regulation of FBL, NHP2L1 and NOP56 by the three UPR arms, we used the subclones of MCF7 cells expressing shRNA targeting PERK, ATF6 and XBP1, respectively. Their knockdown efficiencies were verified by Western blot. Fig 2A-C demonstrates that the PERK, ATF6 and XBP1 expression were effectively reduced in the respective subclones of MCF7 cells. The MCF7 subclones were then treated with or without BFA for 24 h since BFA is a well-established potent UPR activator. However, Fig 2C-D indicates no significant change in the expressions of FBL, NHP2L1 and NOP56 following the knockdown of individual UPR arms. The UPR-induced reduction in the expression of NHP2L1 was compromised in PERK knockdown cells. There is significant cross talk and redundancy among the UPR signaling pathways, and UPR pathways can compensate for each other to some extent, ensuring that the cell can still respond to ER stress even if one pathway is compromised. It is likely that due to redundancy in UPR pathways, downregulation of FBL and NOP56 is not affected when any single UPR arm is impaired in isolation.

**Fig 2.**
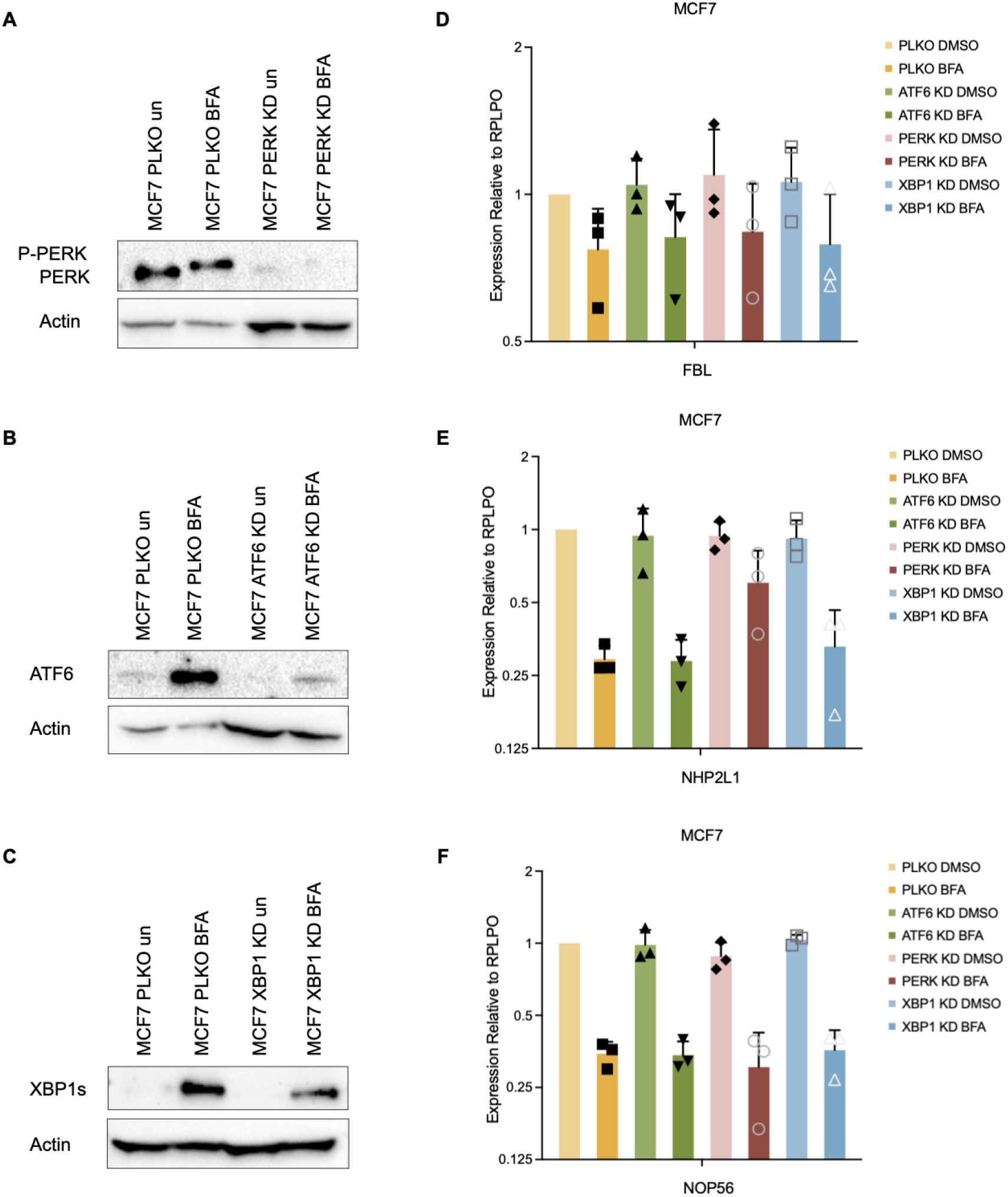
Role of UPR sensors in regulation of FBL, NHP2L1 and NOP56 expression during UPR. Specific shRNAs were used to generate MCF7 knockdown clones for (A) PERK (PERK-KD), (B) ATF6 (ATF6-KD) and (C) XBP1 (XBP1-KD). These clones were treated with either a control or BFA (0.5 μg/ml) for 24 h. Western blot analysis of total protein was performed using antibodies against XBP1s, PERK, ATF6 and β-actin. The expressions of (D) FBL, (E) NHP2L1 and (F) NOP56 were quantified by qPCR (n=3), normalized to RPLPO. Error bars represent mean ±S.D. from three independent experiments conducted in triplicate. Statistical analysis was performed using one-way ANOVA.

### Regulation of rRNA methylation during UPR

Since rRNA half-life is more than 3 days [28], to explore the regulation of rRNA methylation during UPR, the cell viability of MCF7 cells after 96 hours of TG treatment was assessed. Fig 3A shows that the concentration required to achieve ∼50% reduction in cell viability in parental MCF7 cells was near to 0.1 μM after 96 hours of TG treatment. Western blot analysis (Fig 3B and Fig 3C) reveals that the protein levels of FBL and NOP56 were significantly reduced, with a decrease of over 50% and about 30%, respectively. These conditions were used, and RNA was isolated to check methylation of rRNA. The rRNA methylation levels were assessed using the procedure described by Belin *et al*. and is briefly described in the Materials and Methods section [12]. The results demonstrated significant decreases in rRNA methylation levels at positions 1489 of the 18S rRNA and 1858 of the 28S rRNA (Fig 3D). However, based on the RT efficiencies, only the methylation level at position 1489 of the 18S rRNA showed a significant reduction (Fig 3E).

**Fig 3.**
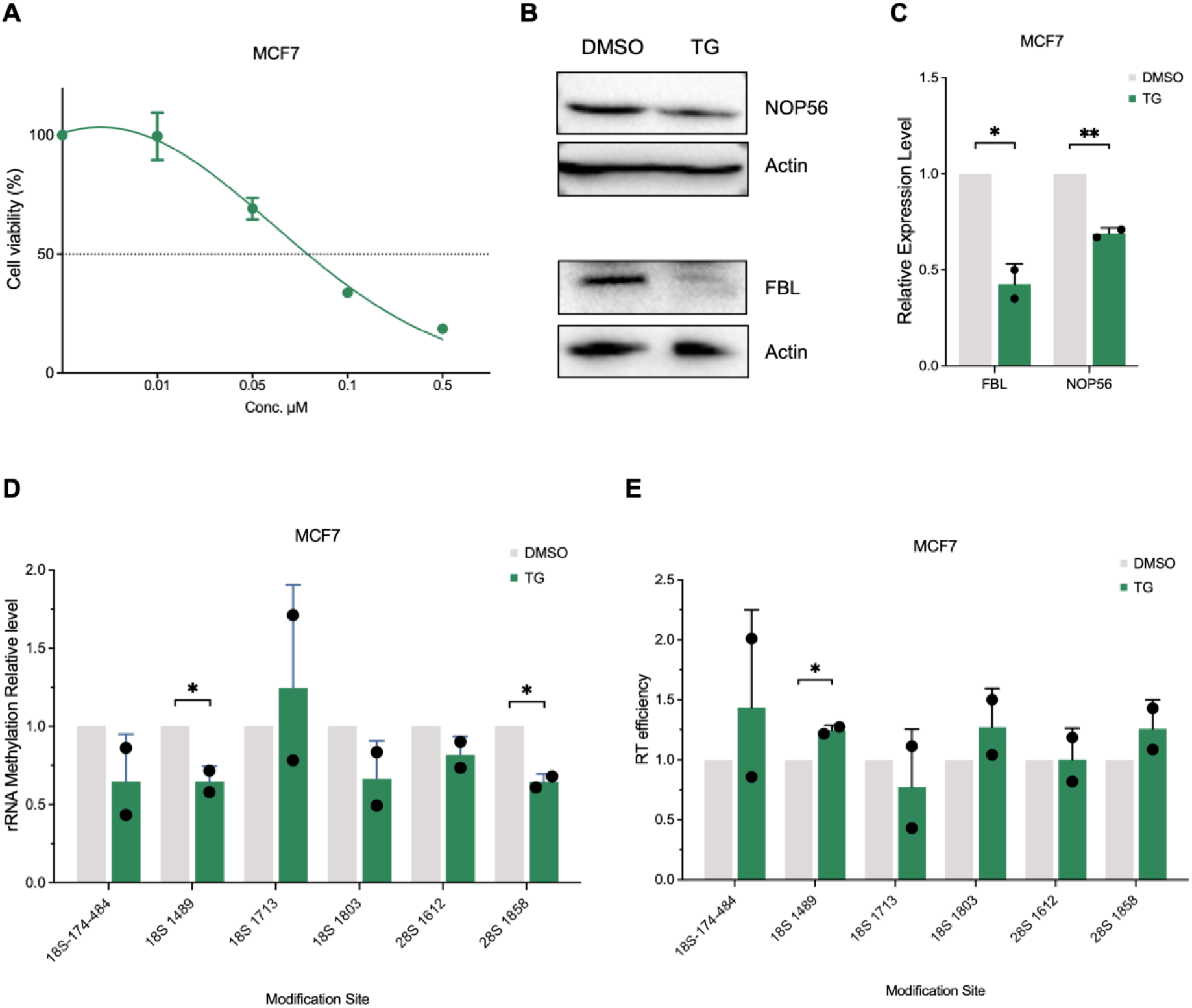
Regulation of rRNA methylation by UPR. (A) Evaluation of cell viability after TG treatment with varying concentrations. Parental MCF7 cells (2,000 per well) were seeded in a 96-well plate and exposed to different TG concentrations for four days. Cell viability was assessed using the MTS assay, measuring absorbance at 490 nm. Error bars represent mean ±S.D. from the data of five replicate wells for each concentration. Statistical analysis was conducted using one-sample t-tests, yielding a significance level of (P=0.0183). (B) Parental MCF7 cells were treated with TG (0.1 μM) for 96 h. Western blot analysis of total protein was conducted using antibodies against NOP56, FBL and β-actin to analyze the expressions of NOP56 and FBL (n=2). The image J quantifications of the blots in (B) were shown in (C). (D, E) Parental MCF7 cells were treated with TG (0.1 μM) for 96 h. The methylation levels were quantified using qPCR (n=2), normalized to RPLPO. The rRNA methylation levels were assessed based on the qPCR data using the methods presented in (D) Ref. [26] and (E) Ref. [27]. Error bars represent mean ±S.D. from the independent experiments. Statistical analysis was performed using unpaired t-tests. *P<0.05 and **P<0.01.

### Regulation of IRES initiated translation and stop codon readthrough during UPR

Ribosomes can act as pivotal regulators in protein translation, affecting cellular function and gene expression by altering their rRNA and protein composition [14]. Six reporter plasmids (Table 1) were used to explore the four crucial ribosomal regulation pathways: ribosome pausing (XBP1u pausing 2A), stop codon readthrough (190UGA), IRES-regulated cap-independent translation and ribosome frameshifting in SARS-COV2. To investigate the role of UPR in translational regulation, parental MCF7 cells were first transfected with four pairs of related plasmids using the Turbofect transfection reagent. These cells were then treated with UPR activators, including BFA and TG. The luciferase reporter assay revealed a significant reduction of approximately 30% in IRES activity following TG treatment in parental MCF7 cells (Fig 4C). The activity of UGA reporter was much lower than activity of WT reporter (three orders on log scale) indicating the stop readthrough occurs a very low frequency. The activity of UGA reporter increased upon treatment with TG and BFA where TG treatment led to robust and significant increase (Fig 4D). There was no effect of UPR on the ribosome pausing and frameshifting activity (Fig 4A and Fig 4B). These results show that UPR can reduce the translation through IRES and increase the stop codon readthrough activity.

**Table 1.** The six plasmids prepared for rRNA regulation.

| Sample Name | Vector size (bp) | Addgene Cat no. |
| --- | --- | --- |
| psiCHECK2-2A-3xFLAG XBP1u_pause-2A | 5698 | 159583 |
| psiCHECK2-2A-3xFLAG-SBP-2A | 5698 | 159584 |
| pFLuc190UGA | 4884 | 41046 |
| pFLucWT | 4884 | 41045 |
| pcDNA3 RLUC POLIRES FLUC | 8836 | 45642 |
| pFRLuc-CoV2-WT | 6501 | 177614 |

**Fig 4.**
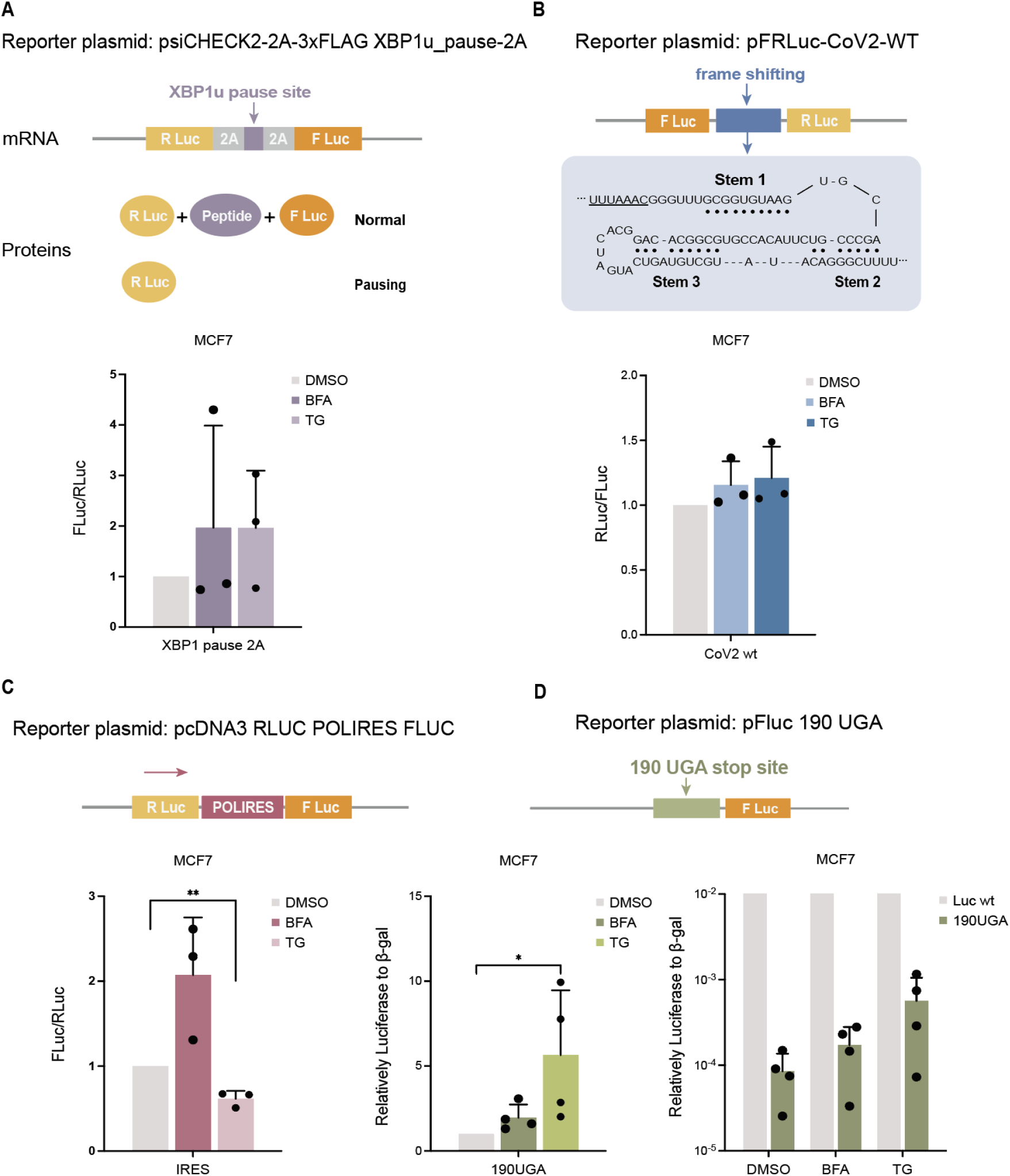
Regulation of translational fidelity during UPR. (A-D) The plasmids were transfected in parental MCF7 cells for 24 h. The cells were then treated with BFA and TG for 24 h. (A-C) The Fluc/Rluc ratio and (D) the Luc/β-gal radio were quantified using reporter assays. Cells were treated with BFA (0.5 µg/ml), TG (1 µM), or DMSO for 24 h post-transfection for the UPR pathway. (A) Transfected with XBP1u pause 2A or vector SBP2A. (B) Transfected with CoV2 WT or untransfected. (C) Transfected with POLIRES or untransfected. (D) Transfected with 190 UGA or vector Luc WT. Error bars represent mean ±S.D. from three or four independent experiments. Statistical analysis was performed using unpaired t-test. *P< 0.05 and **P<0.01.

### Regulation of UPR signaling by FBL

To investigate the role of FBL in rRNA methylation, the MCF7 shFBL subclone was generated. FBL expression was knocked down using tetracycline-inducible shRNA lentivirus. The subclones were treated with doxycycline to activate the Tet-on shRNA system, which silenced FBL by inducing shRNA expression. The qPCR results reveal that the MCF7 shFBL subclones treated with doxycycline demonstrated reductions in FBL expression by 80.4% (Fig 5A). Western blot analysis reveals a time-dependent reduction of FBL expression in MCF7 shFBL following doxycycline treatment (Fig 5B and Fig 5C). Next, the role of FBL in the proliferation and colony formation of breast cancer cells were investigated. The proliferation of MCF7 shFBL and MCF7 scramble subclones began to display significant growth differences starting from the third day (Fig 5D), indicating that FBL knockdown significantly slowed the cell growth rate. MCF7 shFBL subclones also formed fewer colonies after two weeks of culture (Fig 5E and Fig 5F). Notably, although the number of colonies did not significantly decrease when seeding 2000 cells per well, the size of the colonies was much smaller compared to those in MCF7 scramble cells. Further investigations were conducted on how FBL knockdown regulates UPR-related proteins including XBP1s, P-eIF2α, total eIF2α, PERK and CHOP in MCF7 cells. BFA was used to activate UPR pathway for different time points (0, 4 and 8 h) in MCF7 scramble and shFBL subclones. As shown in Fig 5G, no difference in UPR protein expression between MCF7 scramble and MCF7 shFBL cells was observed, with the notable exception P-eIF2α and CHOP. These results suggest that FBL knockdown did not affect the ATF6 or IRE1-XBP1 pathway but suppressed activation of PERK pathway in response to ER stress.

**Fig 5.**
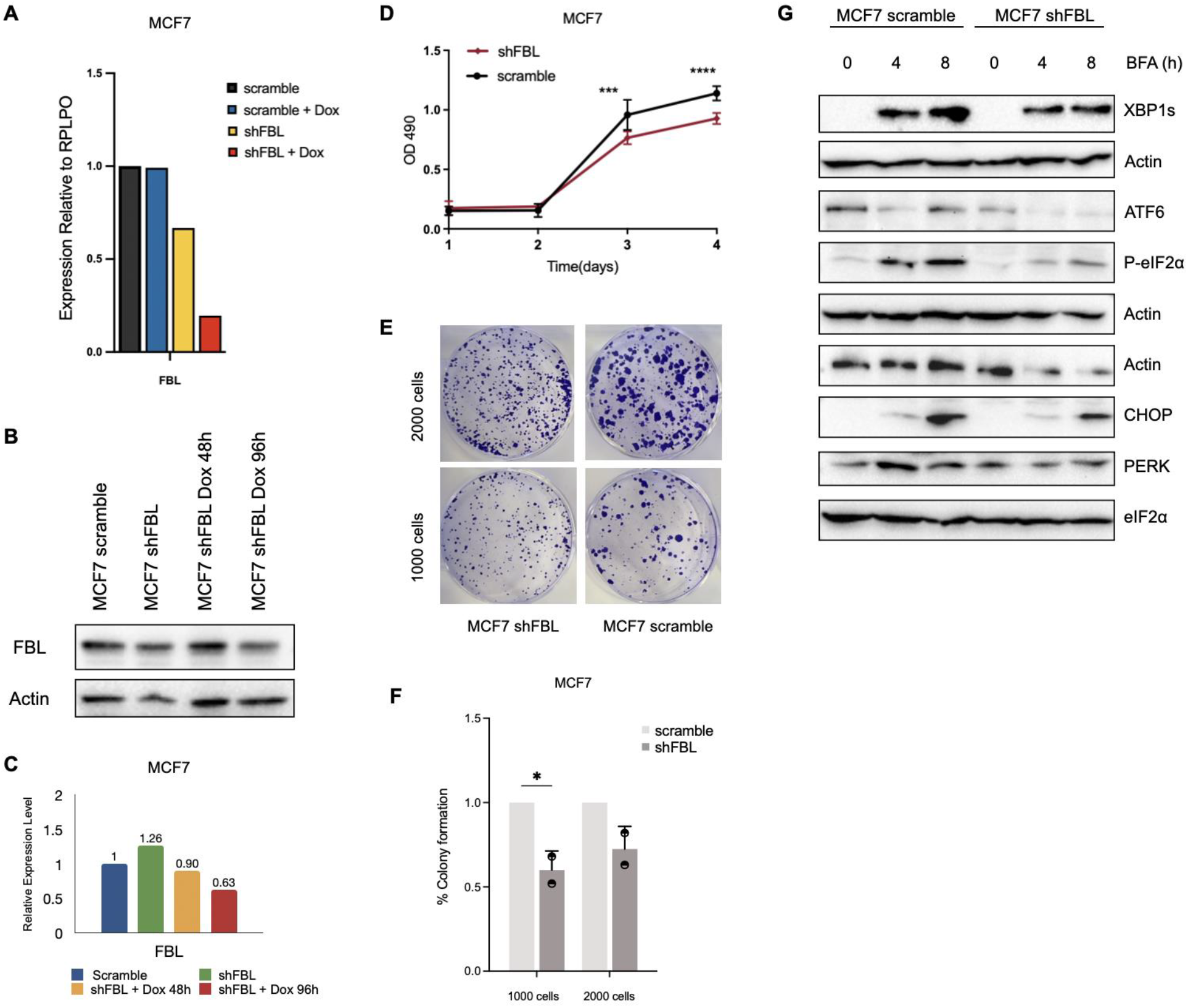
Effect of FBL on cell growth and UPR signaling. (A) The subclones were treated with doxycycline (1 ng/ml) for 48 h. FBL expression was quantified using qPCR, normalized to RPLPO. (B, C) The subclones were treated with doxycycline (1 ng/ml) for 48 h and 96 h. Western blot analysis of total protein was performed using antibodies against FBL and β-actin. The expression of FBL assessed analyzed by Western blot. The image J quantification of the blots in (B) are shown in (C). (D) The subclones were treated with doxycycline (1 ng/ml) for 48 h and seeded into 96-well plates (2,000 cells per well). The cell proliferation was assessed using the MTS assay, measuring absorbance at 490 nm. Data from one of two independent experiments with five replicate wells each are presented as the mean ±S.D. (E, F) The subclones (1,000 or 2,000 per well) were seeded in a 6-well plate. The MCF7 and T47D subclones were cultured for two and three weeks, respectively. The fresh medium with doxycycline (1 ng/ml) was changed every four days. The cells were then fixed with 4% formalin and stain with 0.1% crystal violet. Error bars represent mean ±S.D. from the two independent experiments. Statistical analysis was performed using two-way ANOVA. *P<0.05, ***P<0.001 and ****P<0.0001. (G) The subclones were treated with BFA (0.5 μg/ml) for different time points (0, 4, 8 h) after doxycycline treatment for 48 h. Western blot analysis of total protein was performed using antibodies against XBP1s, eIF2α, PERK, CHOP, P-eIF2α and β-actin.

**Fig 6.**
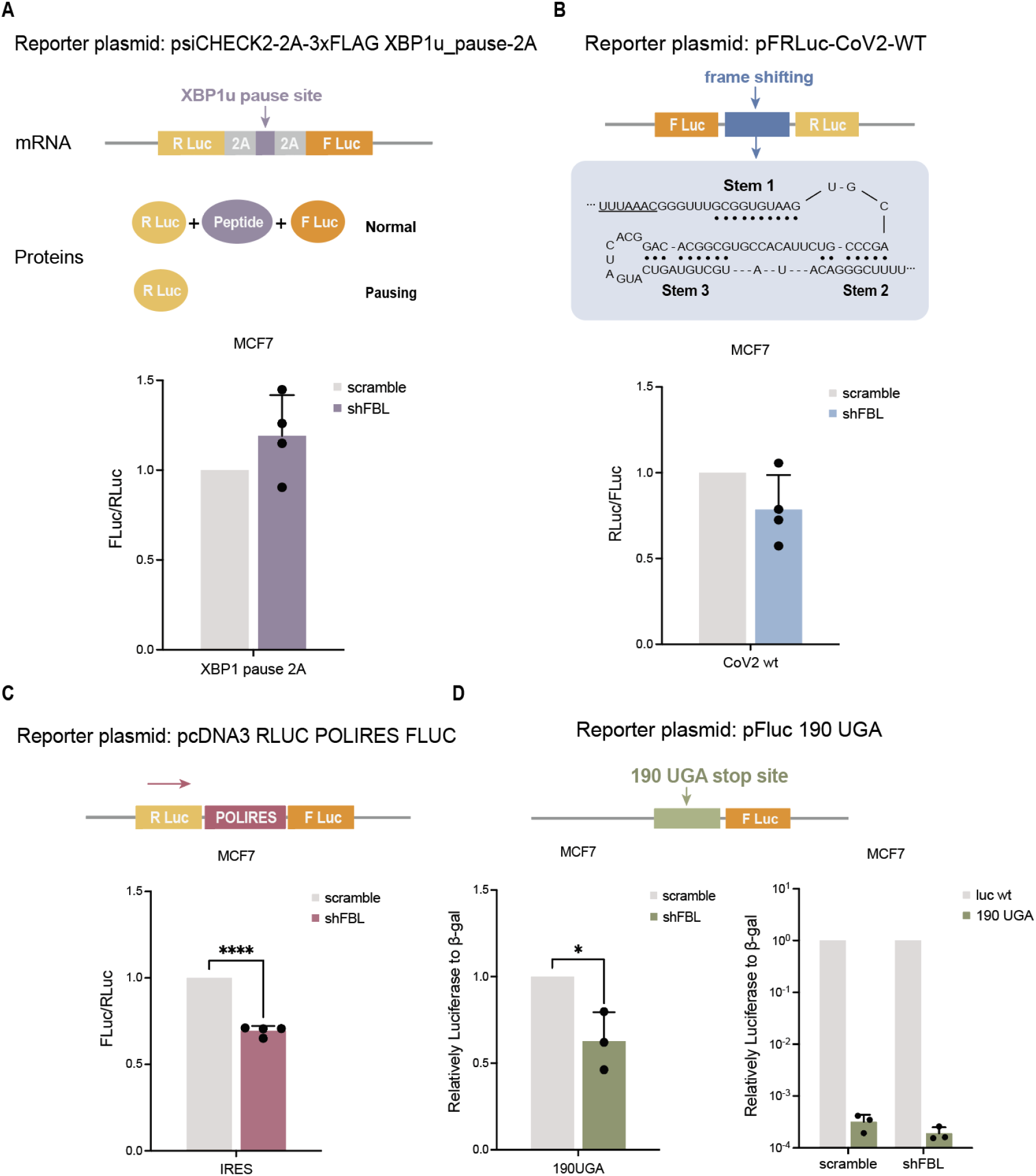
Effect of FBL on translational fidelity. (A-D) MCF7 cells (scramble and shFBL) were treated with doxycycline for 48 h, followed by transfection with specific constructs for 24 h. (A-C) The Fluc/Rluc ratio and (D) the Luc/β-gal radio were quantified using reporter assays. (A) Transfected with XBP1u pause 2A or vector SBP2A. (B) Transfected with CoV2 WT or untransfected. (C) Transfected with POLIRES or untransfected. (D) Transfected with 190 UGA or vector Luc WT. Error bars represent mean ±S.D. from three or four independent experiments. Statistical analysis was performed using unpaired t-test. *P< 0.05 and ****P<0.0001.

### Regulation of IRES initiated translation and stop codon readthrough by FBL

To assess the effect of FBL knockdown on fidelity of translation, four pairs of plasmids were transfected into MCF7 shFBL and scramble subclones. These cells were treated with doxycycline (1 ng/ml) for 48 h in advance. The Luciferase reporter assay revealed that FBL knockdown decreased the activity of IRES reporter (30%) and 190 UGA reporter (35%) (Fig 7). These data suggest that FBL knockdown may fine-tune translation through modulation of IRES and stop codon readthrough pathways.

**Fig 7.**
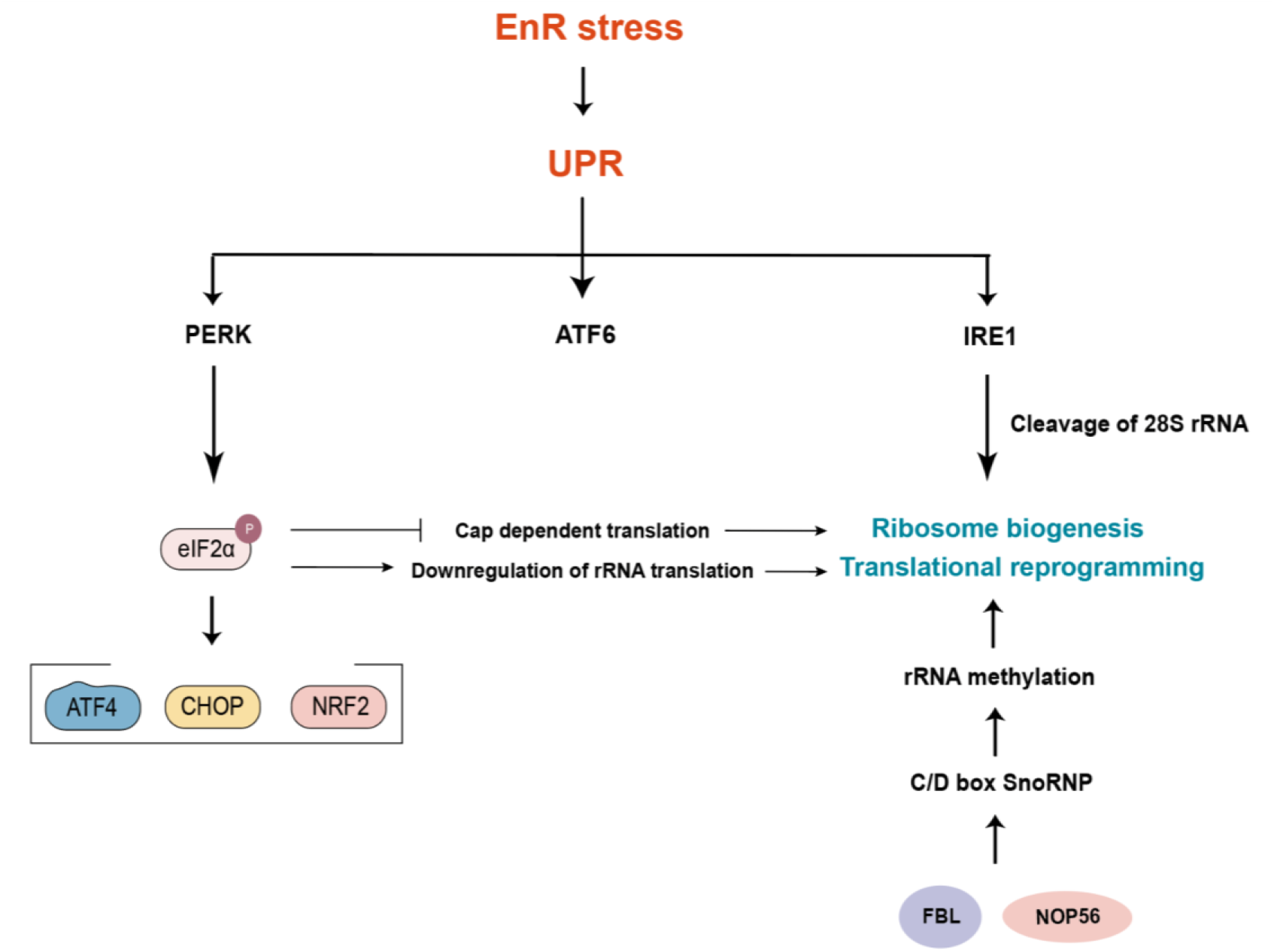
Summary of results. ER stress triggers the UPR pathways, including the ATF6, PERK and IRE1 arms. The PERK arm upregulates P-eIF2α along with its downstream proteins ATF4, CHOP and NRF2. FBL and NOP56 promote rRNA methylation, enhancing ribosome biogenesis and translational reprogramming. P-eIF2α also downregulates rRNA translation and suppresses cap-dependent translation, further facilitating ribosome biogenesis and translational reprogramming. Additionally, IRE1 also affects ribosome biogenesis and translational reprogramming by promoting the cleavage of 28S rRNA.

## Conclusion

Since SnoRNAs were discovered in 1982, the understanding of their functions has evolved from the assumption that their activity was limited to the nucleolus and thus overlooked, to a growing consensus that SnoRNAs are dysregulated in cancer, with differential expression across cancer types, stages and metastasis [29].

The results are summarized as in Fig 8. The qPCR results showed that in both parental MCF7 and T47D cells, NOP56 expression was downregulated after TG and BFA treatments, whereas FBL expression was downregulated after TG treatment. Using Western blot, we verified that the protein levels of NOP56 and FBL were significantly reduced after 96 hours of TG (0.1 μM) treatment. Then we used this condition to check the rRNA methylation levels using qPCR. The results suggest that the rRNA methylation levels at positions 1489 of the 18S rRNA and 1858 of the 28S rRNA were downregulated. Iwawki *et al*. found that IRE1 influenced translation through the cleavage of 28S rRNA under ER stress [30]. Pierce *et al*. provided evidence that IRE1 recognized unspliced HAC1 mRNA (functional homolog of XBP1) associated with ribosomes during ER stress and suggested that ribosome integrity was essential for IRE1 to process HAC1 mRNA [31].

The luciferase reporter assay then revealed that after TG treatment in parental MCF7 cells, IRES activity was significantly reduced, and the readthrough translation via the stop codon 190 UGA increased. MTS and colony formation assays demonstrated that FBL knockdown reduced the proliferation of MCF7 cells. It has been reported that eIF2α is phosphorylated by PERK under ER stress, thereby enhancing the translation of some specific transcription factors, such as ATF4, CHOP and NRF2 [32]. In addition, although the PERK/P-eIF2α pathway initially suppresses protein synthesis to alleviate ER stress, the subsequent activation of ATF4 and its downstream effectors, such as mTORC1, reactivates protein synthesis at levels that support cell survival under chronic stress [33]. We found that FBL knockdown attenuated the UPR-induced upregulation of P-eIF2α and CHOP but had no significant effect on XBP1s and ATF6. These results indicate that FBL knockdown selectively impairs the activation of PERK pathway in response to ER stress without affecting ATF6 and IRE1-XBP1 signaling. The luciferase reporter assay showed that FBL knockdown in MCF7 cells decreased IRES activity and suppressed the readthrough translation via the stop codon 190 UGA. The previous qPCR results have demonstrated that the activation of UPR by TG leads to decreased FBL expression in parental MCF7 and T47D cells. Therefore, we hypothesize that TG may target FBL to suppress the IRES-dependent translation and the readthrough translation through the stop codon 190 UGA. The relationship between FBL and the UPR pathways requires further investigation. Moreover, FBL is related to DNA damage. For example, the FBL-IRES-p53-DNA damage pathway has been reported [34]. Additionally, the FBL knockdown also suppressed the IRES pathway. From this perspective, we can explore how FBL regulates cellular damage through the modulation of IRES following TG treatment in the future.

In conclusion, we found that the expressions of NOP56 and FBL were downregulated during UPR. During the UPR or FBL knockdown, the rRNA methylation levels were altered. Additionally, the stop codon readthrough translation was enhanced during UPR, whereas the IRES-dependent translation was suppressed. However, stop codon readthrough and IRES-dependent translation were suppressed by FBL knockdown.

## Abbreviations

ATF6: activating transcription factor 6
BFA: brefeldin A
BSA: bovine Serum Albumin
CHOP: C/EBP homologous protein
DKC1: dyskerin dyskeratosis congenita 1
DMEM: Dulbecco’s modified eagle’s medium
DPBS: Dulbecco’s phosphate buffered saline
eIF2α: eukaryotic initiation factor 2 alpha subunit
ER: endoplasmic reticulum
FBL: fibrillarin
FBS: fetal bovine serum
GAR1: guided anti-RNP 1
IRE1: inositol requiring enzyme 1
IRES: internal ribosome entry site
KD: knockdown;
KM: Kaplan-Meier
NHP2L1: NHP2-like protein 1
NOP10: nucleolar protein 10
NOP56: nucleolar protein 56
NOP58: nucleolar protein 58
PBST: phosphate buffered saline with tween-20
P-eIF2α: phospho-eukaryotic initiation factor 2α
PERK: protein kinase RNA-like endoplasmic reticulum kinase
PMS: phenazine methosulfate
POLIRES: poliovirus internal ribosome entry site
qPCR: quantitative real-time polymerase chain reaction
RPLPO: ribosomal protein, large,
PO; RQ: relative quantification
rRNA: ribosomal RNA
RT: reverse transcription
SARS-COV2: severe acute respiratory syndrome coronavirus 2
SBP2A: selenocysteine insertion sequence-binding protein 2A
SDS-PAGE: sodium dodecyl sulphate-polyacrylamide gel electrophoresis
SnoRNA: small nucleolar RNA
SNoRNPs: small nucleolar ribonucleoprotein
TBST: tris-buffered saline with tween-20
TBT: tris buffer with tween-20
TG: thapsigargin
TM: tunicamycin
UPR: unfolded protein response
WT: wild type
XBP1: X-box binding protein 1.

## Acknowledgements

We acknowledge the support received from the technical officers of Lambe Institute for Translational Research, University of Galway. W.Z. received a PhD fellowship from the China Scholarship Council (CSC) [grant number 202006370066].

## Author contributions

W.Z., A.G. and S.G. wrote the manuscript. W.Z. and A.G. collected the literature. S.G. primarily revised and finalized the manuscript. W.Z. and A.G. made the figures and revised the manuscript for clarity and style. All authors have read and approved the final manuscript.

## Supporting information

**Table S1.**
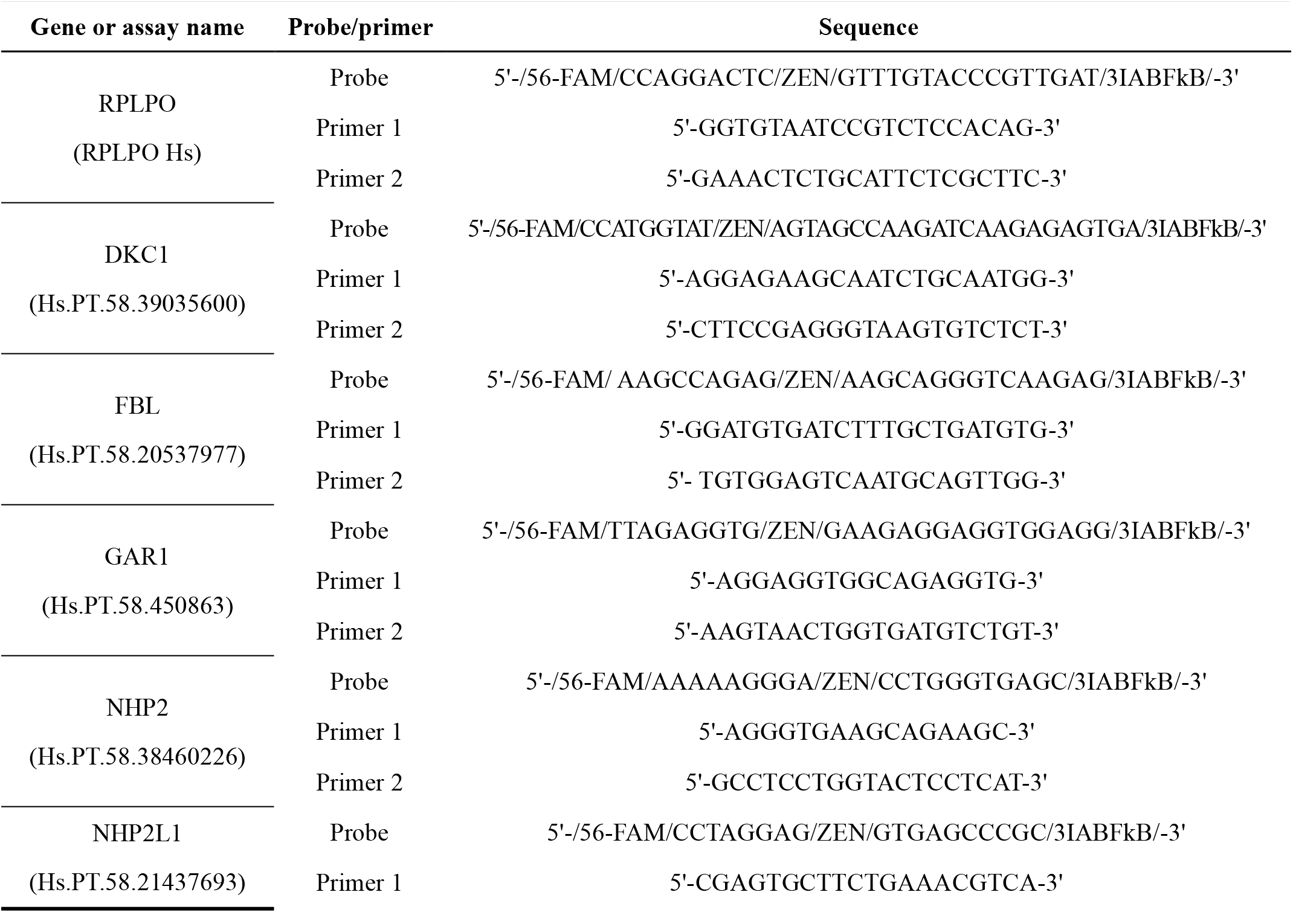
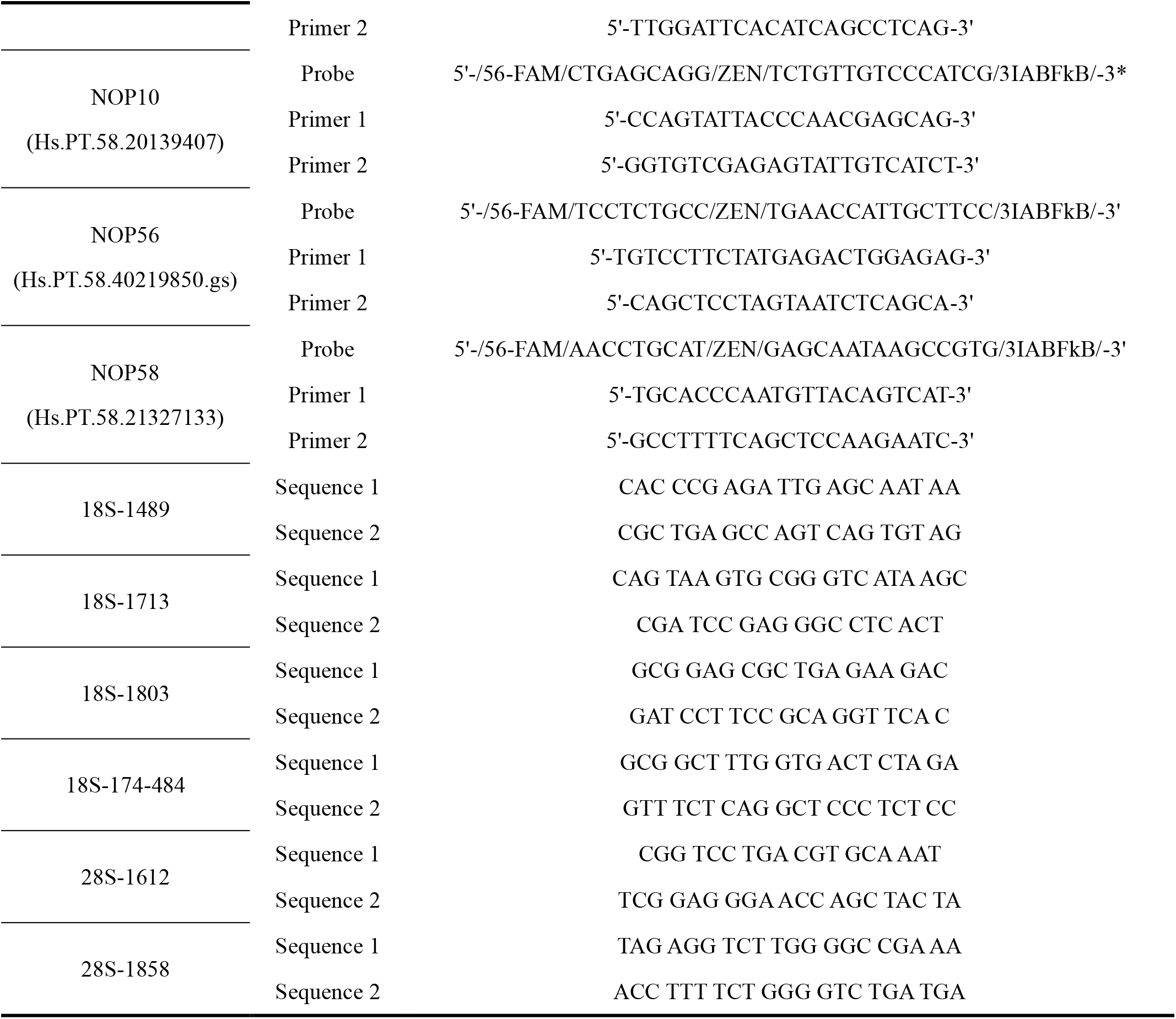
List of PCR primers used to test the expression of genes.

**Table S2.** List of antibodies.

| Ab name | Size (KD) | Cat no. | Source | Blocking (2 h) | Primary incubation (Overnight at 4°C) | Secondary incubation (2 h at RM) |
| --- | --- | --- | --- | --- | --- | --- |
| β-actin | 42 | Sigma- | Rabbit | 5% Milk in | 1:5000 in 5% Milk | Anti-Rabbit Ab |
|  |  | Aldrich |  | PBS/0.05% | in PBS/0.05% | (1:5000) 5% Milk in |
|  |  | A5060 |  | Tween | Tween | PBS/0.05% Tween |
| FBL | 33 | Abcam | Rabbit | 5% BSA in | 1:5000 in 5% Milk | Anti-Rabbit Ab |
|  |  | ab5821 |  | TBS/0.05% | in TBS/0.05% | (1:5000) 5% BSA in |
|  |  |  |  | Tween | Tween | TBS/0.05% Tween |
| NOP56 | 66 | Abcam<br>Ab229497 | Mouse | 5% Milk in<br>PBS/0.05%<br>Tween | 1:1000 in 5% Milk<br>in PBS/0.05%<br>Tween | Anti-Mouse Ab<br>(1:5000) 5% Milk in<br>PBS/0.05% Tween |
| PERK | 140 | Cell Signaling<br>C33E10 | Rabbit | 5% BSA in<br>TBS/0.05%<br>Tween | 1:2000 in 5% BSA<br>in TBS/0.1% Tween | Anti-Rabbit Ab<br>(1:5000) 5% BSA in<br>TBS/0.05% Tween |
| XBP1s | 55 | Bio-legend<br>143F | Mouse | 5% Milk in<br>PBS/0.05%<br>Tween | 1:1000 in 5% Milk<br>in PBS/0.05%<br>Tween | Anti-Mouse Ab<br>(1:5000) 5% Milk in<br>PBS/0.05% Tween |
| CHOP | 30 | Cell Signaling<br>L63F7 | Mouse | 5% Milk in<br>PBS/0.05%<br>Tween | 1:2000 in 5% Milk<br>in PBS/0.05%<br>Tween | Anti-Mouse Ab<br>(1:5000) 5% Milk in<br>PBS/0.05% Tween |
| eIF2α | 38 | Cell Signaling<br>9722S | Rabbit | 5% BSA in<br>TBS/0.05%<br>Tween | 1:2000 in 5% BSA<br>in TBS/0.1% Tween | Anti-Rabbit Ab<br>(1:5000) 5% BSA in<br>TBS/0.05% Tween |
| P-eIF2α | 38 | Cell Signaling<br>9721S | Rabbit | 1% BSA in<br>TBS/0.05%<br>Tween | 1:1000 in 5% BSA<br>in TBS/0.1% Tween | Anti-Rabbit Ab<br>(1:5000) 5% BSA in<br>TBS/0.05% Tween |

